# *Alkaliphilus flagellata* sp. nov., *Butyricicoccus intestinisimiae* sp. nov., *Clostridium mobile* sp. nov., *Clostridium simiarum* sp. nov., *Dysosmobacter acutus* sp. nov., *Paenibacillus brevis* sp. nov., *Peptoniphilus ovalis* sp. nov., and *Tissierella simiarum* sp. nov., isolated from monkey feces

**DOI:** 10.1101/2021.09.23.461610

**Authors:** Dan-Hua Li, Rexiding Abuduaini, Meng-Xuan Du, Yu-Jing Wang, Hong-He Chen, Nan Zhou, Hai-Zhen Zhu, Yong Lu, Pei-Jun Yu, Shu-Peng Yang, Cheng-Ying Jiang, Qiang Sun, Chang Liu, Shuang-Jiang Liu

**Author notes:** Corresponding authors: Shuang-Jiang Liu (or) & Chang Liu, Postal address: Institute of Microbiology, Chinese Academy of Sciences, Beichen West Road No. 1, Chaoyang District, Beijing 100101, China. Author contributions: Conceived and designed the experiments: Shuang-Jiang Liu, Chang Liu, Qiang Sun. Performed the experiments: Dan-Hua Li, Rexiding Abuduaini, Meng-Xuan Du, Yu-Jing Wang, Yong Lu, Pei-Jun Yu, Shu-Peng Yang. Analyzed the data: Dan-Hua Li, Chang Liu, Hai-Zhen Zhu, Hong-He Chen, Zhou Nan, Cheng-Ying Jiang. Drafted the manuscript: Dan-Hua Li, Chang Liu. Approved final version of manuscript: Shuang-Jiang Liu. All authors read and approved the final manuscript. **The NCBI/NMDC accession numbers of the 16S rRNA gene and genome sequences for the 8 type strains are**: (i) MSJ-1^T^, 16S rRNA sequence: MZ310594 and NMDCN0000NQV; genome sequence: JAHLQO000000000/ NMDC60018343; (ii) MSJ-2^T^, 16S rRNA sequence: MZ310595/NMDCN0000NR0; genome sequence: JAHLQN000000000/NMDC60018344; (iii) MSJ-4^T^, 16S rRNA sequence: MZ310597/NMDCN0000NR2; genome sequence: JAHLQL000000000/NMDC60018346; (iv) MSJ-5^T^, 16S rRNA sequence: MZ310598/NMDCN0000NR3; genome sequence: JAHLQK000000000/NMDC60018347; (v) MSJ-6^T^, 16S rRNA sequence: MZ310599/NMDCN0000NR4; genome sequence: JAHLQJ000000000/NMDC60018348; (vi) MSJd-7^T^, 16S rRNA sequence: MZ310600/NMDCN0000NR5; genome sequence: JAHLQI000000000/NMDC60018349; (vii) MSJ-11^T^, 16S rRNA sequence: MZ310603/NMDCN0000NR8; genome sequence: JAHLQF000000000/NMDC60018352; (viii) MSJ-40^T^, 16S rRNA sequence: MZ310625/NMDCN0000NRU; genome sequence: JAHLPM000000000/NMDC60018374.

## Abstract

Non-human primates harbor diverse microbiomes in their guts. As a part of China Microbiome Initiatives, we cultivated and characterized the gut microbiome of cynomolgus monkeys (*Macaca fascicularis*). In this report, we communicate the characterization and taxonomy of 8 bacterial strains that were obtained from fecal samples of captive cynomolgus monkeys. The results revealed that they represented 8 novel bacterial species. The proposed names of the 8 novel species are *Alkaliphilus flagellate* (type strain MSJ-5^T^ =CGMCC 1.45007^T^=KCTC 15974^T^), *Butyricicoccus intestinisimiae* MSJd-7^T^ (type strain MSJd-7^T^ =CGMCC 1.45013^T^ =KCTC 25112^T^), *Clostridium mobile* (type strain MSJ-11^T^ =CGMCC 1.45009^T^=KCTC 25065^T^), *Clostridium simiarum* (type strain MSJ-4^T^ =CGMCC 1.45006^T^ =KCTC 15975^T^), *Dysosmobacter acutus* (type strain MSJ-2^T^ =CGMCC 1.32896^T^=KCTC 15976^T^), *Paenibacillus brevis* MSJ-6^T^ (type strain MSJ-6^T^ =CGMCC 1.45008^T^=KCTC 15973^T^), *Peptoniphilus ovalis* (type strain MSJ-1^T^ =CGMCC 1.31770^T^=KCTC 15977^T^), and *Tissierella simiarum* (type strain MSJ-40^T^ =CGMCC 1.45012^T^=KCTC 25071^T^).

## Introduction

The gastrointestinal tracts accommodate diverse microbes, and those microbes together in a host gastrointestinal tract are called gut microbiomes (GMs) [1]. Many efforts have been made to characterize the microbial diversities of human [2–6] and animal GMs [7, 8], by culture-dependent and/or -independent methods [9–11]. Based on analysis of major and large-scale investigations of human gut microbiomes [2–6], there are 5,000-6,000 bacterial species associated with human beings. Exploration of cultivated human gut bacterial species diversity revealed that about 1,500 bacterial species were recorded with standings of valid and correct names, and that more gut bacterial species were cultivated but not characterized or nominated [2]. Those cultivated but unnamed bacterial species remained as “uncultivated” in the databases server such as “Silva” [12] or “GTDB” [13], and they would be repeatedly claimed as “novel bacteria” by later-on studies. Bacterial cultivation, characterization with polyphasic methods, and deposits in culture collections are essential to nominate a bacterial name following the rules of International Code of Nomenclature of Prokaryotes (ICNP). Considering the large numbers (usually more than thousands) of bacterial isolates from one microbiome study, it is a challenge to characterize and nominate all bacterial isolates from microbiome studies.

Nonhuman primates (NHPs) are the most biologically relevant animal models for human studies [14]. The compositions and dynamics of NHPs gut microbiomes were evaluated using culture-independent methods [15–18]. In addition, members of the genera *Bacteroides*, *Bifidobacterium*, *Eubacterium*, *Fusobacterium*, *Lactobacillus*, and *Streptococcus* were cultivated and reported from the gastrointestinal tract of NHPs [19]. The China Microbiome Initiatives (CMI) integrated multiple efforts on studies of human and animal gut microbiomes and environmental microbiomes [20]. As a part of CMI, we cultivated and characterized the gut microbiome of cynomolgus monkeys (*Macaca fascicularis*). In this report, we present the characterization and taxonomy of 9 bacterial strains that were obtained from fecal samples of captive cynomolgus monkeys.

## Materials and methods

### Sample collection and treatment

All fecal samples were from cynomolgus monkeys (*M. fascicularis*) at the experimental animal center of Institute of Neuroscience, Chinese Academy of Sciences, Suzhou, China. Fresh fecal samples were collected and maintained in airtight bags on dry ice, and were delivered immediately to the laboratory The samples were diluted with sterile PBS (phosphate buffered solution) and filtered through a 40 μm cell strainer and were treated with 70 % ethanol or heated at 85 °C for 30 min, as described in literatures [21, 22].

### Culture media, bacterial isolation and cultivation

The following media were used for bacterial cultivation: FAB medium (Solarbio, LA4550) and YCFA medium [23], modified GAM medium (mGAM) [2, 24], and modified R medium [25]. The modified GAM medium (per 1 L) contained 10 g casitone, 3 g soya peptone, 15 g proteose peptone, 13.5 g digested serum, 5 g yeast extract, 2 g beef extract powder, 1.2 g liver extract, 0.3 g soluble starch, 0.5 g L-cysteine, 0.5 g L-arginine, 0.3 g L-tryptophan, 2 g NaHCO_3_, 2.5 g KH_2_PO_4_, 3 g NaCl, 0.15 g CH_2_(SH)COONa, 2.46 g CH_3_COONa, 0.01 g hemin, 0.001 g resazurin, 0.3 g glucose, 0.3 g D-galactose, 0.3 g D-cellobiose, 0.3 g mannose, 0.3 g fructose, 0.3 g rhamnose, 0.3 g palatinose, 0.3 g inulin, 15 g agar, adjusted pH to 7.2, sterilized at 115 °C for 25 min. The modified R medium was prepared from two solutions that were prepared and sterilized separated: Solution A (per 900 mL) consisted of 6 g casein hydrolysate, 5 g peptone, 5 g yeast extract, 1 g glucose, 1 g inulin, 1 g D-fructose, 1 g D-cellobiose, 1.5 g NaCl, 0.1 g MgSO4. H_2_O, 5 mL hemin (0.1 %, w/v), 1 mL resazurin (0.1 %, w/v), 20 mL (2 %, v/v) rumen fluid, 15 g agar, adjusted pH to 7.2, sterilized at 112 °C for 15 min. Solution B (per 100 mL) consisted of 0.4 g L-cysteine, 1 g ascorbic acid, 0.1 g glutathione, 2 g α-ketoglutarate, 0.45 g K_2_HPO_4_, 0.9 g KH_2_PO_4_, adjusted pH to 7.2, filtered using a 0.2 μm micro filter.

Sterilized agar plates were inoculated with dilutions of pretreated fecal samples and incubated at 37 °C under strictly anaerobic conditions with N_2_ (85 %), H_2_ (10 %) and CO_2_ (5 %) in an anaerobic chamber (Electrotek AW400SG workstation, West Yorkshire, UK). Colonies appeared after cultivation for 2, 5 and 10 days were picked and re-streaked on agar plates of same media. Bacterial purity was evaluated by observation of morphology, 16S rRNA gene and genome sequencing.

### Cell morphology observation, chemotaxonomic determinations

Cell morphology was determined by transmission electron microscopy (JEM-1400; JEOL). The utilization of carbon sources was determined using the 96 well BIOLOG AN microplate (BIOLOG Inc., Hayward, CA, USA) that contained 95 different carbon substrates [26]. Bacteria strains were cultured in liquid mGAM medium for 2 days, then cells were harvested. Cellular fatty acids were extracted and methylated according to the standard protocol of MIDI (Sherlock Microbial Identification System, version 6.0). The identification was performed by GC (HP 6890 Series GC System; Agilent) [27]. Polar lipids were separated by two-dimensional thin layer chromatography (TLC plates coated with silica gel, 1010 cm; Merck). Chromatography was performed using chloroform/methanol/water (65 : 25 : 4, in volume) for the first dimension, followed by chloroform/methanol/acetic acid/water (80 : 12 : 15 : 4, in volume) for the second dimension [28]. Total lipids were detected with 10 % ethanolic molybdatophosphoric acid (Sigma). Aminolipids were detected with 0.4 % solution of ninhydrin (Sigma) in butanol. Phospholipids were detected with Zinzadze reagent (molybdenum blue spray reagent, 1.3 %; Sigma) and glycolipids were detected with 0.5 % α-naphthol reagent.

### Fermentative production of Short-chain fatty acids

Bacterial strains were cultivated for 72 h in mGAM broth at 37 °C under strictly anaerobic conditions. Short-chain fatty acids (SCFAs) were measured using GC-MS method. 1 mL culture was extracted with 1 mL ethyl acetate. The supernatant liquid was prepared for GC-MS analysis which was performed on a GCMS-QP2010 Ultra with an auto sampler (SHIMADZU) and the DB-wax capillary column (30 m, 0.25 mm i.d., 0.25 μm film thickness, SHIMADZU). The temperature of oven was programmed from 80 °C to 140 °C at 20 °C/min gradient, with 1 min hold; to 290 °C at 3.5 °C/min, with 15 min hold. Injection of 1 μL sample was performed at 280 °C. The carrier gas, helium, flowed at 1.2 mL/min. Electronic impact was recorded at 70eV [33].

### 16S rRNA gene sequencing and phylogenetic analysis

Complete 16S rRNA gene sequences of isolates were obtained using the universal primers 27F (5’-AGAGTTTGATCCTGGCTCAG-3’) and 1492R (5’-GGTTACCTTGTTACGACTT-3’). 16S rRNA gene sequences similarities were determined using EzBioCloud server [29]. Multiple alignments of sequences were performed using the CLUSTAL W [30]. The phylogenetic trees were constructed by the neighbor-joining (NJ) method [31] according to Kimura’s two-parameter model [32] in MEGA X [33], and also by maximum-likelihood (ML) method [34] based on the Tamura-Nei model and maximum parsimony (MP) method [35] based on Subtree-Pruning-Regrafting (SPR) search method. The statistical reliability of the trees was calculated by bootstrap analysis with 1000 replications [36].

### Genome sequencing and analysis

Genomic DNA was extracted using the Wizard Genomic DNA Purification Kit (Promega) and genomic DNA library was sequenced on an Illumina Hiseq X-ten platform. All good quality paired reads were assembled using the SPAdes software (v3.9.0) [37]. The up-to-date bacterial core gene set UBCG (https://www.ezbiocloud.net/tools/ubcg) [38] was used to extract closely related and available genomes and to generate genome-based phylogenomic tree. The average nucleotide identity (ANI) value of the closely related and available genomes was calculated using OAT software at http://www.ezbiocloud.net/sw/oat along with UPGMA dendrogram (unweighted pair group method with arithmetic mean) [39]. The genomic distances, digital DNA-DNA hybridization (dDDH), were calculated by using the Genome-To-Genome Distance Calculator (GGDC; http://ggdc.dsmz.de/) [40]. The percentage of conserved proteins (POCP) was calculated by the method previously described [41]. Genome analysis using the Check M indicated that the genomes of eight strains were not contaminated [42].

### Culture preservations

Bacterial strains were cultured in liquid medium for 2 days. We stored our own cultures (1 mL) in lab by addition of equal volume of 65 % (v/v,) glycerol (1 mL), and was placed at −80°C for long-term preservation. All type strains assigned by this study were deposited at China General Microbiological Culture Collection Center (CGMCC) and Korean Collection for Type Culture (KCTC), and strain numbers are included in the section of species description.

## Results and Discussion

### Source and isolation of the bacteria

Strains MSJ-1^T^, MSJ-2^T^, MSJ-4^T^, MSJ-5^T^, MSJ-6^T^, MSJd-7^T^, MSJ-11^T^ and MSJ-40^T^ were isolated from feces samples of *Macaca fascicularis*. Strain MSJ-1^T^ was obtained from sample after enrichment, strain MSJ-4^T^ and MSJ-5^T^ were obtained from samples after heat treatment (85 °C for 30 min), and strains MSJ-2^T^, MSJ-6^T^, MSJd-7^T^, MSJ-11^T^ and MSJ-40^T^ were obtained from samples after 70 % ethanol treatment for 30 min. These strains MSJ-1^T^/MSJ-2^T^/MSJ-4^T^/MSJ-11^T^ were successfully cultivated first with FAB, strain MSJ-5^T^ first from YCFA, strain MSJd-7^T^ first from modified mGAM, and MSJ-40^T^ first from modified R media. But we demonstrated later that they all grew with mGAM medium.

### Bacterial growth and cell morphology

Strains MSJ-1^T^, MSJ-2^T^, MSJ-4^T^, MSJ-5^T^, MSJ-6^T^, MSJd-7^T^, MSJ-11^T^ and MSJ-40^T^ were strictly anaerobic bacteria. They grew on mGAM agar at 37 °C and formed visible colonies after 1-10 days. Colonies were white or grey, no pigments were observed, and additional features are provided in the species description. Cellular morphology was examined with transmission electron microscopy and are showed in Fig. 1. Strains MSJ-1^T^ and MSJd-7^T^ were spherical-shaped. MSJ-2^T^, MSJ-4^T^, MSJ-5^T^, MSJ-6^T^, MSJ-11^T^ and MSJ-40^T^ were rod-shaped. Flagella were observed for strains MSJ-4^T^, MSJ-5^T^, MSJ-6^T^, MSJ-11^T^, and MSJ-40^T^ but not for MSJ-1^T^, and MSJd-7^T^. Additional features of those strains are detailed as in the species description.

### Assimilation of carbon sources and fermentative production of short chain fatty acids

The assimilation of 95 carbon sources were tested with BIOLOG AN microplates and results are recorded in Fig. 2a. The 8 bacteria showed different carbon source spectrum, and totally 75 out of the 95 carbon sources were metabolized. We found that mono- and di-saccharides were preferred by the strains, which extensively exist in guts [43]. The 8 bacteria assimilated in common for five carbon sources, i.e., D-fructose, L-fucose, D-galacturonic acid, palatinose and pyruvic acid.

Many gut microbes produce short chain fatty acids that are related host health [43, 44]. We determined the production of short chain fatty acid in mGAM broth that contained glucose, D-galactose, D-cellobiose, mannose, fructose, rhamnose, palatinose, and inulin. The results showed that each strain produced unique profiles of short chain fatty acids (Fig. 2b). Butyric acid was produced by MSJ-1^T^, MSJ-4^T^, MSJ-5^T^, MSJ-6^T^, MSJ-11^T^ and MSJ-40^T^. Propionic acid was produced MSJ-4^T^, MSJ-11^T^, MSJ-6^T^ and MSJ-40^T^. Acetic acid was produced by MSJ-1^T^, MSJ-2^T^, MSJ-4^T^, MSJ-5^T^, MSJ-6^T^, MSJ-11^T^ and MSJ-40^T^. In addition to the above SCFAs, strains MSJ-6^T^ and MSJ-11^T^ produced also branched SCFAs of isobutyric acid and/or isovaleric acid. The strain MSJd-7^T^ did not produce the six SCFAs detected in this study.

### Cellular fatty acid and polar lipid profiling

Chemotaxonomic features of cellular fatty acids and polar lipid profiles for the 8 bacteria were determined and are summarized. As showed in Table 1, the 8 bacteria had different cellular fatty acid profiles, but they all had C_14:0_, C_16:0_, C_18:0_ and anteiso _C15:0_. Taking 10 % as cutoff value for predominant cellular fatty acids, MSJ-1^T^ had C_16:0_ (19.92 %), MSJ-2^T^ had C_16:0_ (20.05 %), MSJ-4^T^ had C_14:0_ (15.16 %) and C_16:0_ (24.48 %), MSJ-5^T^ had C_16:0_ (10.62 %) and iso-C_13:0_ (11.25 %)/anteiso-C_17:0_ (14.5 %)/anteiso-C_15:0_ (15.74 %)/iso-C_16:0_ (16.79 %), MSJ-6^T^ had C_16:0_ (20.59 %)/C_18:0_ (10.47 %) and iso-C_16:0_ (17.12 %)/anteiso-C_15:0_ (25.63 %), MSJd-7^T^ had C_14:0_ (10.93 %)/C_16:0_ (26.75 %)/C_18:0_ (21.95 %) and iso-C_17:1 ω5c_ (12.84 %), MSJ-11^T^ had C_14:0_ (19.8 %)/C_16:0_ (37.37 %)/C_18:0_ (11.67 %), and MSJ-40^T^ had mainly iso-C15:0 (62.15 %). Polar lipid profiling showed that all 8 bacteria had diphosphatidylglycerol (DPG) and phosphatidylglycerol (PG), but were different from each other in the presence or not of phosphatidylethanolamine (PE), phosphatidylmethylethanolamine (PME), unknown phospholipids (PL) and unknown lipids (L), unknown glycolipids, as detailed in Table 1 and supplementary Fig. S1.

**Table 1.**
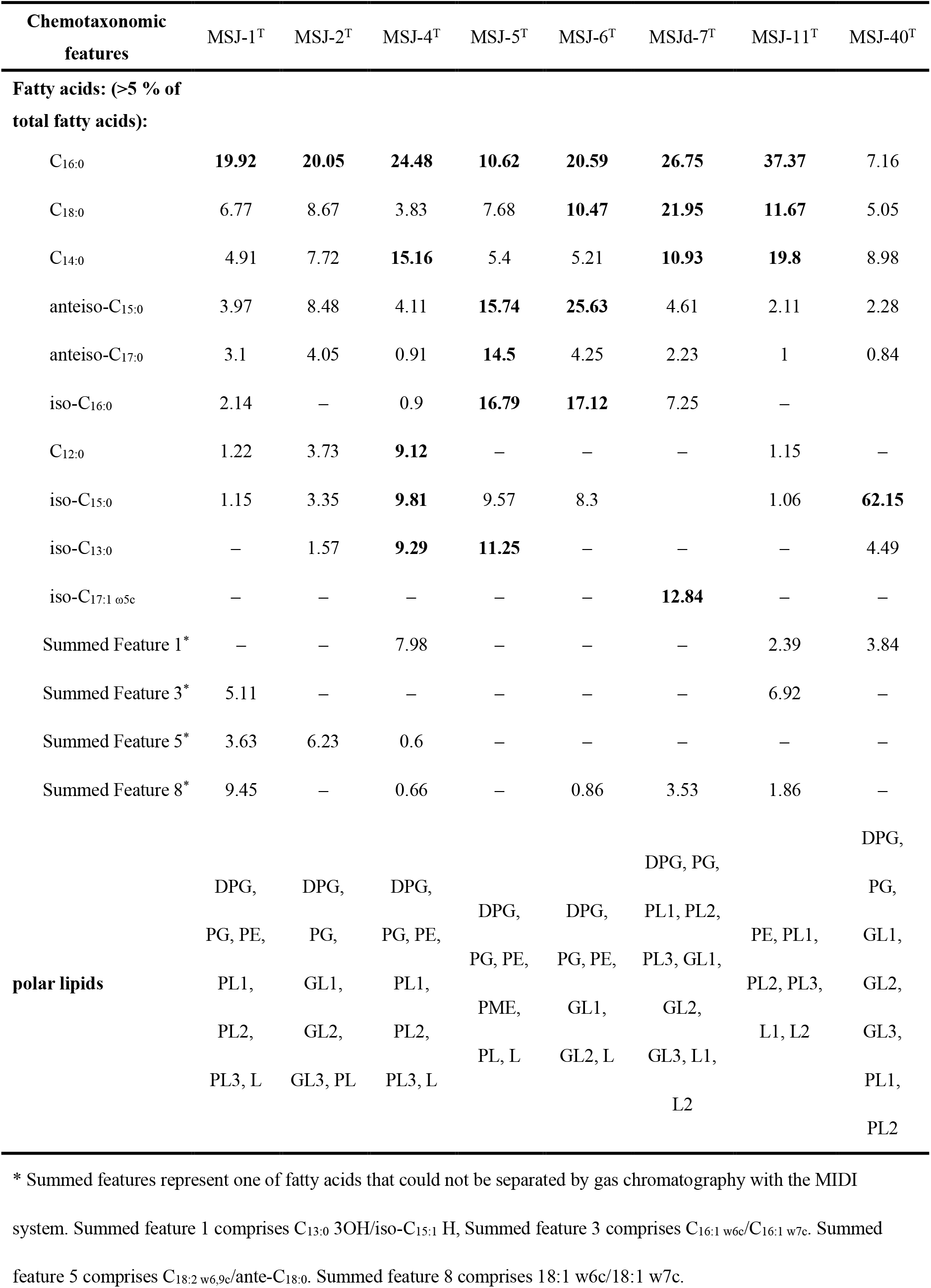
Chemotaxonomic and general genome features of 8 bacteria

### General features of genome and genomic DNA G+C contents

The 8 bacteria had different genome sizes, MSJ-1^T^ was the smallest (2.1 Mbp) and MSJ-6 ^T^ was the largest genome (5.2 Mbp). Genome sequencing data and some basic features coding density and G+C contents are listed in Table 2. The genomic DNA G+C molar contents of MSJ-1^T^, MSJ-2^T^, MSJ-4^T^, MSJ-5^T^, MSJ-6^T^, MSJd-7^T^, MSJ-11^T^ and MSJ-40^T^ were 30.65 %, 58.27 %, 30.46 %, 31.71 %, 49.3 %, 50.29 %, 30.38 %, 44.49 % and 30.39 %, respectively.

**Table 2.** Genome features of eight strains

| Genome features | MSJ-1<sup>T</sup> | MSJ-2<sup>T</sup> | MSJ-4<sup>T</sup> | MSJ-5<sup>T</sup> | MSJ-6<sup>T</sup> | MSJd-7<sup>T</sup> | MSJ-11<sup>T</sup> | MSJ-40<sup>T</sup> |
| --- | --- | --- | --- | --- | --- | --- | --- | --- |
| Genome Size (bp) | 2102036 | 3161374 | 3811517 | 3614516 | 5239947 | 2711934 | 4014245 | 4088863 |
| G+C content<br>(mol %) | 30.65 | 58.27 | 30.46 | 31.71 | 49.3 | 50.29 | 30.38 | 30.39 |
| Completeness | 98.6 | 94.63 | 99.19 | 97.87 | 98.66 | 99.33 | 100 | 99.13 |
| Contamination | 0.7 | 0.67 | 0.93 | 0.24 | 0 | 0.67 | 0.57 | 1.98 |
| Number of contigs | 14 | 2 | 70 | 51 | 64 | 27 | 18 | 53 |
| N50 of contigs (bp) | 300523 | 3156307 | 406816 | 534169 | 198601 | 357991 | 1137742 | 199075 |
| Gene number | 2037 | 3296 | 3532 | 3591 | 4800 | 3023 | 4017 | 4177 |

### The 8 bacterial strains represent novel taxa

Based on the 16S rRNA gene and genomic data, we further investigated the phylogenetic and phylogenomic relationships of the 8 bacteria to their closely related and currently validly nominated bacterial taxa (Fig. 3a, Fig. S2, Fig. S3 and Fig. 3b). ANI scores were used to generated UPGMA dendrogram trees (Fig. 3c). Combining the results from DNA molecule analysis and the phenotypic characterization, we concluded that MSJ-1^T^, MSJ-2^T^, MSJ-4^T^, MSJ-5^T^, MSJ-6^T^, MSJd-7^T^, MSJ-11^T^, and MSJ-40^T^ represented novel species of the currently known genera (for details, see following paragraphs).

#### Strain MSJ-1^T^

The phylogenetic trees revealed that strain MSJ-1^T^ clustered within the previously described genus *Peptoniphilus* clade, supported by 100 % bootstrap value (Fig. 3a). Strain MSJ-1^T^ was closest related to *Peptoniphilus gorbachii* WAL10418^T^ (96.82 % identity), *Peptoniphilus lacydonensis* DSM 100661^T^ (95.93 %) and *Peptoniphilus harei* DSM 10020^T^ (95.64 %) [45–47]. In phylogenomic tree, strain MSJ-1^T^ also formed a separate branch located in the genus *Peptoniphilus* clade (Fig. 3b). ANI and dDDH between strain MSJ-1^T^ and the closest neighbor *P. lacydonensis* DSM 100661^T^ (GCA 900106515.1) were 78.79 % and 20.50 %, respectively (Fig. 3c). At the time of writing, the genus *Peptoniphilus* contains 20 species with validly published names [48]. Cells of *Peptoniphilus* members are non-spore-forming, obligately anaerobic and coccus-shaped. In addition to the unique 16S rRNA gene and genome sequence, MSJ-1^T^ contains the major fatty acid of C_16:0_ that is consistent with the most species of the genus *Peptoniphilus*, although the fatty acids of other members in the genus *Peptoniphilus* have more diversity [49]. Based on phenotypic, chemotaxonomic, phylogenetic analysis, phylogenomic tree and genome data, we suggest that strain MSJ-1^T^ represents a novel species affiliated to the genus *Peptoniphilus* and the name *Peptoniphilus ovalis* sp. nov. is proposed.

#### DESCRIPTION OF *PEPTONIPHILUS OVALIS* SP. NOV

*Peptoniphilus ovalis* sp. nov. (o.va’lis. L. fem. adj. *ovalis*, pertaining to an egg, egg-shaped). Cells are non-mobile cocci with the diameter of approximately 0.6-0.8 μm, and no flagellum. Strictly anaerobic, heterotrophic growth at temperature and pH are 37 °C and 7.0, respectively. Produces butyric acid and acetic acid from fermentation. After 2 days of cultivation on mGAM agar plate, colonies were 1-2 mm in diameter, white, circular, entire, opaque and smooth. The strain MSJ-1^T^ metabolizes dextrin, D-fructose, L-fucose, D-galactose, D-galacturonic acid, gentiobiose, α-D-glucose, glucose-6-phosphate, lactulose, D-mannose, D-melibiose, 3-methyl-D-glucose, palatinose, L-rhamnose, glyoxylic acid, α-hydroxybutyric, ß-hydroxybutyric, α-ketobutyric acid, α-ketovaleric acid, D- and L-lactic acid, D-lactic acid methyl ester, D-malic acid, pyruvic acid, pyruvic acid methyl ester, urocanic acid, L-alanyl-L-histidine, L-glutamic acid, L-glutamine, L-serine, 2’-deoxy adenosine, inosine, thymidine, uridine, thymidine-5’-monophosphate and uridine-5’-monophosphate. The predominant cellular fatty acid (>10 %) is C_16:0_ (19.92 %). The polar lipids are diphosphatidylglycerol, phosphatidylglycerol, phosphatidylethanolamine, three unknown phospholipids and an unknown lipid. Genome size is 2,102,036 bp and the G+C content is 30.65 mol %. The name of *Peptoniphilus ovalis* sp. nov. is proposed. The type strain is MSJ-1^T^ (=CGMCC 1.31770^T^=KCTC 15977^T^) and was isolated from fecal samples of *M. fascicularis*.

#### Strain MSJ-2^T^

MSJ-2^T^ was closely related to *Dysosmobacter welbionis* DSM 106889^T^ (95.78 % 16S rRNA gene identity) [50] and *Oscillibacter valericigenes* DSM 18026^T^ (95.15 %) and *Oscillibacter ruminantium* JCM 18333^T^ (94.72 %) [51, 52]. Phylogenetic tree viewing indicated MSJ-2^T^ formed a heterogeneous cluster of members of *Dysosmobacter* and *Oscillibacter* (Fig. 3a and 3b). The OrthoANI tree clearly separated MSJ-2^T^ and *D. welbionis* from *O. valericigenes* and *O. ruminantium* (Fig. 3c). Furthermore, the G+C molar content of strain MSJ-2^T^ (58.27 %) was closer to that of *D. welbionis* (58.9 %) than to *O. valericigenes* (53.2 %) and *O. ruminantium* (55.0 %). Thus, MSJ-2^T^ was more likely a member of genus *Dysosmobacter*. ANI and dDDH between strain MSJ-2^T^ and the closest cultivated neighbor *D. welbionis* DSM 106889^T^ (GCA 005121165.1) were 74.09 % and 21.20 %, respectively, suggesting they were different species within the genus *Dysosmobacter*. Cells of strain MSJ-2^T^ were long rods, with no flagella. The predominate fatty acid of MSJ-2^T^ is C_16:0_ (20.59 %) that distinguished further this organism from *D. welbionis* DSM 106889^T^ (C_16:0_ < 1 %) [50]. Based on the chemotaxonomic, phylogenetic and genomic analysis described above, we concluded that stain MSJ-2^T^ represented a novel species affiliated to the genus *Dysosmobacter* and the name *Dysosmobacter acutus* sp. nov. is proposed.

#### DESCRIPTION OF *DYSOSMOBACTER ACUTUS* SP. NOV

*Dysosmobacter acutus* sp. nov. (acu.t’u.s N.L. masc. adj. *acutus*, sharp, pointed referring to atypical cell shape). Cells are non-mobile long rods with sharp ends. No flagellum. The cell size is approximately 0.5-0.6 μm x 2.7-2.9 μm. Strictly anaerobic, heterotrophic growth at temperature and pH of 37 °C and 7.0, respectively. Colonies were < 1 mm in diameter after 5 days of incubation at 37 °C on mGAM agar plates, plat, circular, entire, translucent and smooth. Fermentative production of isovaleric acid, isobutyric acid and acetic acid. The strain MSJ-2^T^ metabolizes D-cellobiose, dextrin, D-fructose, L-fucose, D-galactose, D-galacturonic acid, gentiobiose, α-D-glucose, glucose-6-phosphate, lactulose, D-mannose, D-melibiose, 3-methyl-D-glucose, palatinose, L-rhamnose, acetic acid, formic acid, glyoxylic acid, α-ketobutyric acid, propionic acid, pyruvic acid and pyruvic acid methyl ester. The predominant cellular fatty acid (>10 %) is C_16:0_ (20.05 %). The major lipids are phosphatidylglycerol and two unknown glycolipids. Genome size is 3,161,374 bp and the G+C content is 58.27 mol %. The name of *Dysosmobacter acutus* sp. nov. is proposed. The type strain is MSJ-2^T^ (=CGMCC 1.32896^T^=KCTC 15976^T^), which was isolated from a fecal sample of *M. fascicularis*.

#### Strain MSJ-4^T^ and strain MSJ-11^T^

Based on the phylogenetic and phynogenomic trees, strains MSJ-4^T^ and MSJ-11^T^ formed a cluster but well separated to the validly nominated members of genus *Clostridium* [53]. MSJ-4^T^ was closely related to *Clostridium algidicarnis* DSM 15099^T^ (96.85 % 16S rRNA gene identity) and *Clostridium putrefaciens* NCTC 9836^T^ (96.78 %) [54, 55]. MSJ-11^T^ was closely related to *Clostridium malenominatum* ATCC 25776^T^ (98.33 %) [56]. The ANI and dDDH of strain MSJ-4^T^ to its closest neighbor *C. algidicarnis* DSM15099^T^ (GCA 002934235.1) were 78.04 % and 21.60 %, respectively. The ANI and dDDH values of strain MSJ-11^T^ to its closest cultivated neighbor *Clostridium lundense* DSM 17049^T^ (GCA 000619945.1) were 75.24 % and 21.60 %, respectively (Fig. 3a~3c). Our results revealed that the major fatty acids compositions of strain MSJ-4^T^ and MSJ-11^T^ were C_16:0_, C_14:0_ and C_12:0_ that were consistent with the majority of species within genus *Clostridium* [57, 58]. Strain MSJ-4^T^ and MSJ-11^T^ belonged to genus *Clostridium* and could be differentiated from each other and from other species of genus *Clostridium*. Therefore, we concluded that strain MSJ-4^T^ represented a novel species and the name of *Clostridium simiarum* sp. nov. is proposed, and that strain MSJ-11^T^ represented also a novel species and the name of *Clostridium mobile* sp. nov. is proposed.

#### DESCRIPTION OF *CLOSTRIDIUM SIMIARUM* SP. NOV

*Clostridium simiarum* sp. nov. (si.mi.a’rum L. neut. pl. n. *simiarum*, of monkeys). Cells are fat rods with blunt ends, 0.5-0.9 μm × 1.4-2.0 μm, and have peritrichous flagella. Strictly anaerobic, heterotrophic growth at temperature and pH of 37 °C and 7.0. Produce white, flat, circular, entire, opaque, smooth colonies with a diameter of 2-3 mm after 2 days of incubation at 37 °C on mGAM agar plates. Strain produces butyric acid, propanoic acid and acetic acid, isovaleric acid and isobutyric acid during fermentation. Assimilates the following carbon sources: dextrin, D-fructose, L-fucose, D-galacturonic acid, palatinose, acetic acid, formic acid, pyruvic acid and pyruvic acid methyl ester. The predominant cellular fatty acids (>10 %) are C_16:0_ (24.48 %) and C_14:0_ (15.16 %). The polar lipids are diphosphatidylglycerol, phosphatidylglycerol, and phosphatidylethanolamine, three unknown phospholipids and an unidentified lipid. Genome size is 3,811,517 bp and the G+C content is 30.46 mol %. The name of *Clostridium simiarum* sp. nov. is proposed. The type strain is MSJ-4^T^ (=CGMCC 1.45006^T^=KCTC 15975^T^) and was isolated from fecal samples of *M. fascicularis*.

#### DESCRIPTION OF *CLOSTRIDIUM MOBILE* SP. NOV

*Clostridium mobile* sp. nov. (mo’bi.le L. neut. adj. *mobile*, motile). Cells are rods with size of approximately 0.4-0.7 μm × 2.9-9.9 μm, have flagella at both ends. Strictly anaerobic, and growth temperature and pH are 37 °C and 7.0, respectively. Strain MSJ-11^T^ produced grey, convex, circular, entire, opaque colonies with a diameter of 2-3 mm after 2 days of incubation at 37 °C on mGAM agar plates. The SCFAs produced by anaerobic fermentation are butyric acid, propanoic acid and acetic acid. The strain MSJ-11^T^ metabolizes D-cellobiose, dextrin, D-fructose, L-fucose, D-galactose, D-galacturonic acid, gentiobiose, α-D-glucose, glucose-6-phosphate, D-mannose, D-melibiose, 3-methyl-D-glucose, palatinose, L-rhamnose, turanose, glyoxylic acid, α-ketobutyric acid, pyruvic acid and pyruvic acid methyl ester. The predominant cellular fatty acids (>10 %) are C_16:0_ (37.37 %), C_14:0_ (19.8 %) and C_18:0_ (11.67 %). The polar lipids are phosphatidylethanolamine as major lipids, minor lipids included three unknown phospholipids and two unknown lipids. Genome size is 4,014,245 bp and the G+C content is 30.38 mol %. The name of *Clostridium mobile* sp. nov. is proposed. The type strain is MSJ-11^T^ (=CGMCC 1.45009^T^=KCTC 25065^T^) and was isolated from fecal samples of *M. fascicularis*.

#### Strain MSJ-5^T^

Strain MSJ-5^T^ was closely related to *Alkaliphilus halophilus* CGMCC 1.5124^T^ (96.29 % 16S rRNA gene identity) and *Alkaliphilus oremlandii* DSM 21761^T^ (96.14 %) [59, 60]. Phylogenetic and phylogenomic trees revealed that strain MSJ-5^T^ was in the genus *Alkaliphilus* clade. The ANI and dDDH values of strain MSJ-5^T^ to its closest neighbor *A. oremlandii* DSM 21761^T^ (GCA 000018325.1) were 75.27 % and 21.20 %, respectively (Fig. 3a~3c). Strain MSJ-5^T^ had rod-shaped and motile cells, which was consistent with the description of the genus *Alkaliphilus*, and had G+C content of 31.71 %, within the range of 28-36 mol % for genus *Alkaliphilus* [59]. The predominant fatty acids composition varied among *Alkaliphilus* species but iso-C_15:0_, iso-C_13:0_, C_16:0_ and C_14:0_ are the major components in most of species, as strain MSJ-5^T^ had. The presence of anteiso-C_15:0_ and anteiso-C_17:0_ was the characteristics of strain MSJ-5^T^ and distinguish this isolate from other *Alkaliphilus* species. Based on the polyphasic analysis, strain MSJ-5^T^ should be classified as a novel species of the genus *Alkaliphilus* and the name of *Alkaliphilus flagellata* sp. nov. is proposed.

#### DESCRIPTION OF *ALKALIPHILUS FLAGELLATA* SP. NOV

*Alkaliphilus flagellata* sp. nov. (fla.gel.la’ta L. neut. n. *flagellum*, a whip; L. fem. adj. suff. -*ata*, suffix denoting provided with; L. fem. part. adj. *flagellata*, flagellated). Cells are rods, 0.7-1.0 μm × 2.2-4.2 μm, have bundled flagella at both ends. Strictly anaerobic, heterotrophic growth at temperature and pH of 37 °C and 7.0, respectively. Produces grey, low convex, circular, entire, opaque colonies with a diameter of 1-2 mm after 2 days of incubation at 37 °C on mGAM agar plates. Fermentative products were butyric acid and acetic acid. The strain MSJ-5^T^ metabolizes dextrin, D-fructose, L-fucose, D-galactose, D-galacturonic acid, gentiobiose, D-glucosaminic acid, α-D-glucose, glucose-6-phosphate, lactulose, D-mannose, D-melibiose, 3-methyl-D-glucose, palatinose, L-rhamnose, α-ketobutyric acid, α-ketovaleric acid, pyruvic acid, pyruvic acid methyl ester, L-alanyl-L-glutamine, L-glutamic acid, L-glutamine, glycyl-L-glutamine, L-methionine, L-serine, L-threonine, 2’-deoxy adenosine, inosine, thymidine and uridine. The major cellular fatty acids (>10 %) are iso-C_16:0_ (16.79 %), anteiso-C_15:0_ (15.74 %), anteiso-C_17:0_ (14.5 %), iso-C_13:0_ (11.25 %) and C_16:0_ (10.62 %). The polar lipids are diphosphatidylglycerol, phosphatidylglycerol, phosphatidylethanolamine, phosphatidylmethylethanolamine, an unknown phospholipid and an unknown lipid. Genome size of is 3,614,516 bp and the G+C content was 31.71 mol %. The name of *Alkaliphilus flagellata* sp. nov. is proposed. The type strain is MSJ-5^T^ (=CGMCC 1.45007^T^=KCTC 15974^T^) and was isolated from fecal samples of *M. fascicularis*.

#### Strains MSJ-6^T^

Strain MSJ-6^T^ was closely related to *Paenibacillus apis* JCM 31620^T^ [61], with 96.94 % 16S rRNA gene sequence identity. The phylogenetic and phylogenomic analysis showed that strain MSJ-6^T^ was in the genus *Paenibacillus* clade (Fig. 3a~3c). The genome size of MSJ-6^T^ was 5239947 bp. The ANI and dDDH values of strain MSJ-6^T^ to its closest neighbor *Paenibacillus faecis* DSM 23593^T^ (GCA 008084145.1) were 72.99 % and 19.50 %, respectively. The unique 16S rRNA and genome sequence is one of the characteristics of MSJ-6^T^. The predominant cellular fatty acids for *Paenibacillus* species were anteiso-C_15:0_, C_16:0_, iso-C_16:0_ and iso-C_15:0_. Strain MSJ-6^T^ shared this feature of cellular fatty acids but the presence of C_18:0_ differentiate it from other *Paenibacillus* species. In addition to diphosphatidylglycerol, phosphatidylglycerol and phosphatidylethanolamine shared by *Paenibacillus* species, MSJ-6^T^ had two unknown glycolipid (GL1 and GL2). In term of phenotypic, chemotaxonomic, phylogenetic and genomic features, strain MSJ-6^T^ should be classified as a novel species of the genus *Paenibacillus* and the name of *Paenibacillus brevis* sp. nov. is proposed.

#### DESCRIPTION OF *PAENIBACILLUS BREVIS* SP. NOV

*Paenibacillus brevis* sp. nov. (bre’vis L. masc. adj. *brevis*, short, denoting the formation of short rods). Cells are ovoid to short rods with size of approximately 0.3-1.42 μm × 2.57-3.57 μm, have 1-2 flagella. Strictly anaerobic, heterotrophic growth at temperature and pH of 37 °C and 7.0, respectively. After 5 days of incubation at 37 °C on mGAM agar plates, colonies were 1-2 mm in diameter, grey, circular, entire, translucent. The SCFAs produced by anaerobic fermentation are acetic acid, valeric acid, propanoic acid, butyric acid and isobutyric acid. The strain MSJ-6^T^ metabolizes amygdalin, D-cellobiose, α-cyclodextrin, β-cyclodextrin, dextrin, dulcitol, I-erythritol, D-fructose, L-fucose, D-galactose, D-galacturonic acid, gentiobiose, D-glucosaminic acid, α-D-glucose, glucose-1-phosphate, glucose-6-phosphate, m-inositol, α-D-lactose, lactulose, maltose, maltotriose, D-mannitol, D-mannose, D-melezitose, D-melibiose, 3-methyl-D-glucose, α-methyl-D-galactoside, β-methyl-D-galactoside, palatinose, D-raffinose, L-rhamnose, salicin, D-sorbitol, stachyose, sucrose, D-trehalose, turanose, acetic acid, fumaric acid, glyoxylic acid, α-ketobutyric acid, α-ketovaleric acid, propionic acid, pyruvic acid, pyruvic acid methyl ester, urocanic acid, L-alanine, glycyl-L-methionine, L-methionine, L-phenylalanine, L-serine, L-valine and L-valine plus L-aspartic acid. The predominant cellular fatty acids (>10 %) are anteiso-C_15:0_ (25.63 %), C_16:0_ (20.59 %), iso-C_16:0_ (17.12 %) and C_18:0_ (10.47 %). The polar lipids are diphosphatidylglycerol, phosphatidylglycerol, phosphatidylethanolamine, two unknown glycolipids and an unknown lipid. Genome size is 5,239,947 bp and the G+C content is 49.3 mol %. The name of *Paenibacillus brevis* sp. nov. is proposed. The type strain is MSJ-6^T^ (=CGMCC 1.45008 ^T^=KCTC 15973^T^) and was isolated from fecal samples of *M. fascicularis*.

#### Strain MSJd-7^T^

Strain MSJd-7^T^ was closely related to *Butyricicoccus porcorum* ATCC TSD-102^T^ [62], with 97.15 % 16S rRNA gene sequence identity. The genome size of MSJd-7^T^ is 2711934 bp. The phylogenetic and phylogenomic analysis revealed that strain MSJd-7^T^ was a member of the genus *Butyricicoccus* clade (Fig. 3a~3c). The ANI and dDDH values of strain MSJd-7^T^ to its closest neighbor *B. porcorum* ATCC TSD-102^T^ (GCA 002157465.1) were 75.88 % and 21.10 %, respectively. *Butyricicoccus* species had diverse cellular fatty acid compositions, but the major components are C_14:0_, C_18:0_ and C_16:0_ in most species of the genus *Butyricicoccus*. The presence of iso-C_17:1 ω5c_ is the characteristic of MSJd-7^T^ and distinguished MSJd-7^T^ from *Butyricicoccus* species. According to the phenotypic, chemotaxonomic, phylogenetic and genomic features, strain MSJd-7^T^ should be classified as a novel species of the genus *Butyricicoccus* and the name of *Butyricicoccus intestinisimiae* sp. nov. is proposed.

#### DESCRIPTION OF *BUTYRICICOCCUS INTESTINISIMIAE* SP. NOV

*Butyricicoccus intestinisimiae* sp. nov. (in.tes.ti’ni si’mi.ae L. masc. n. *intestini*, of the intestine; L. mas. n. *simiae*, of a monkey; N.L.masc. n. *intestinisimiae*, of the monkey intestine, where the type strain dwells). Cells are cocci with a diameter of approximately 1.46-1.85 μm, and no flagellum. Strictly anaerobic, heterotrophic growth at temperature and pH of 37 °C and 7.0, respectively. Colonies were 2-3 mm in diameter after 2 days of incubation at 37 °C on mGAM agar plates, white, convex, circular, entire, opaque and smooth. Anaerobic and fermentative production of valeric acid. The strain MSJd-7^T^ metabolizes i-erythritol, D-fructose, L-fucose, D-galactose, D-galacturonic acid, gentiobiose, D-glucosaminic acid, α-D-glucose, glucose-6-phosphate, D-mannose, D-melibiose, 3-methyl-D-glucose, palatinose, L-rhamnose, D-malic acid, pyruvic acid, succinamic acid, succinic acid and succinic acid Mono-methyl ester. The predominant cellular fatty acids (>10 %) are C_16:0_ (26.75 %), C_18:0_ (21.95 %), iso-C_17:1_ ω5c (12.84 %) and C_14:0_ (10.93 %). The polar lipids are diphosphatidylglycerol, phosphatidylglycerol, four unknown phospholipids, three unknown glycolipids, and two unknown lipids. Genome size is 2,711,934 bp and the G+C content is 50.29 mol %. The name of *Butyricicoccus intestinisimiae* sp. nov. is proposed. The type strain is MSJd-7^T^ (=CGMCC 1.45013=KCTC 25112) and was isolated from fecal samples of *M. fascicularis*.

#### Strain MSJ-40^T^

Strain MSJ-40^T^ was closely related to *Tissierella carlieri* DSM 23816^T^ (94.2 % 16S rRNA gene sequence identity), *Tissierella praeacuta* DSM 18095^T^ (94.13 %) and *Tissierella pigra* DSM 105185^T^ (92.9 %) [63–65]. According to the phylogenetic and phylogenomic trees (Fig. 3a and 3b), MSJ-40^T^ clustered with members of genus *Tissierella*. Thus, strain MSJ-40^T^ was likely be a member of the genus *Tissierella*. As previously reported for *Tissierella* species [64], cells of strain MSJ-40^T^ were rod-shaped. The ANI and dDDH values of strain MSJ-40^T^ to its closest related neighbor *T. pigra* DSM 105185^T^ (GCA 009695605.1) were 74.61 % and 22.40 % (Fig. 3c). The predominant fatty acid of MSJ-40^T^ was iso-C_15:0_, which is a characteristic of genus *Tissierella* [64,65]. At the time of writing, genus *Tissierella* has 5 described species with validly published names, and strain MSJ-40^T^ is different from them according to phenotypic, chemotaxonomic, phylogenetic and genomic features. Thus, strain MSJ-40^T^ should be classified as a novel species of the genus *Tissierella* and the name of *Tissierella simiarum* sp. nov. is proposed.

#### DESCRIPTION OF *TISSIERELLA SIMIARUM* SP. NOV

*Tissierella simiarum* **sp. nov**. (si.mi.a’rum L. neut. pl. n. *simiarum*, of monkeys). Cells are rod-shaped with size of approximately 0.6-0.8 μm × 1.0-3.3 μm, have flagella at both ends. Strictly anaerobic, heterotrophic growth at temperature and pH of 37 °C and 7.0, respectively. The SCFAs produced by fermentation are isovaleric acid, butyric acid, isobutyric acid, propanoic acid and acetic acid. The strain MSJ-40^T^ metabolizes D-cellobiose, dextrin, D-fructose, L-fucose, D-galactose, D-galacturonic acid, gentiobiose, α-D-glucose, glucose-6-phosphate, lactulose, D-mannose, D-melibiose, 3-methyl-D-glucose, palatinose, L-rhamnose, glyoxylic acid, α-ketobutyric acid, pyruvic acid and pyruvic acid methyl ester. The predominant cellular fatty acids (>10 %) are iso-C_15:0_ (62.15 %). The polar lipids are diphosphatidylglycerol and phosphatidylglycerol, three unknown glycolipids and two unknown phospholipids. Genome size is 4.09 Mb and the G+C content is 30.4 mol %. The name of *Tissierella simiarum* sp. nov. is proposed. The type strain is MSJ-40^T^ (=CGMCC 1.45012^T^ =KCTC 25071^T^) and was isolated from fecal samples of *M. fascicularis*.

## Acknowledgements

We thank Prof. Yuguang Zhou at Institute of Microbiology, Chinese Academy of Sciences (CAS) for coordination of deposits of type strains.

