## Supplementary Material for "*Alkaliphilus flagellata* sp. nov., *Butyricicoccus intestinisimiae* sp. nov., *Clostridium mobile* sp. nov., *Clostridium simiarum* sp. nov., *Dysosmobacter acutus* sp. nov., *Paenibacillus brevis* sp. nov., *Peptoniphilus ovalis* sp. nov., and *Tissierella simiarum* sp. nov., isolated from monkey"

*Corresponding authors:

Telephone number: +86-010-64807423

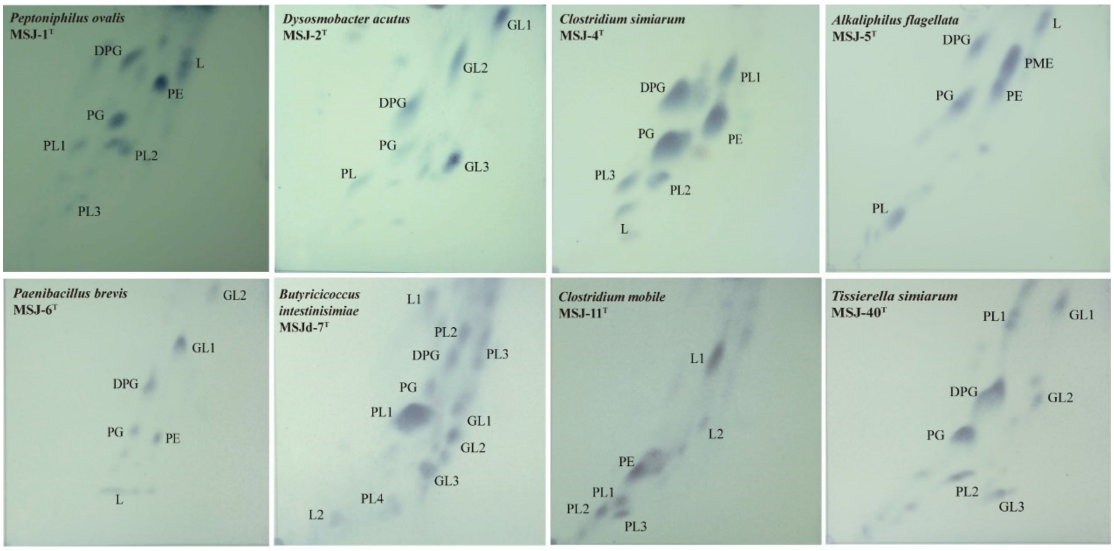


**Fig. S1 Polar lipid profiles after separation by two-dimensional thin layer chromatography of nine isolated strains.**

DPG, diphosphatidylglycerol; PG, phosphatidylglycerol; LPG, Lysophosphatidyl glycerol. PE, phosphatidylethanolamine; PME, phosphatidylmethylethanolamine.

PL, phospholipids; PN, aminophospholipid; GL, glycolipids.

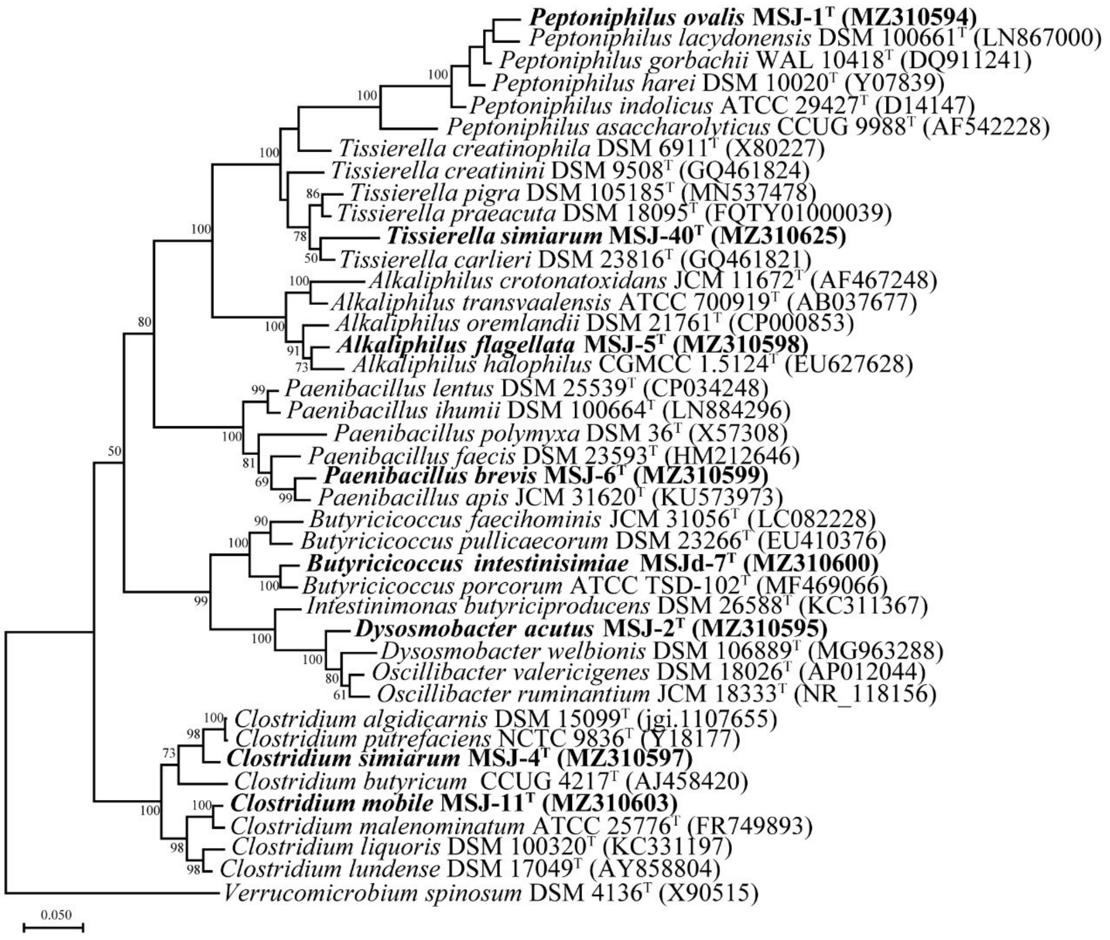


**Fig. S2 Maximum-likelihood phylogenetic tree based on 16S rRNA gene sequences shows the relationship between eight strains and closely related microorganisms.**

Bootstrap percentages (>50%) based on 1,000 replicates are shown at the nodes. GenBank accession numbers are given in parentheses. *Verrucomicrobium spinosum* DSM 4136$^{T}$ (X90515) was used as outgroup. Bar, 0.05 substitutions per nucleotide position.

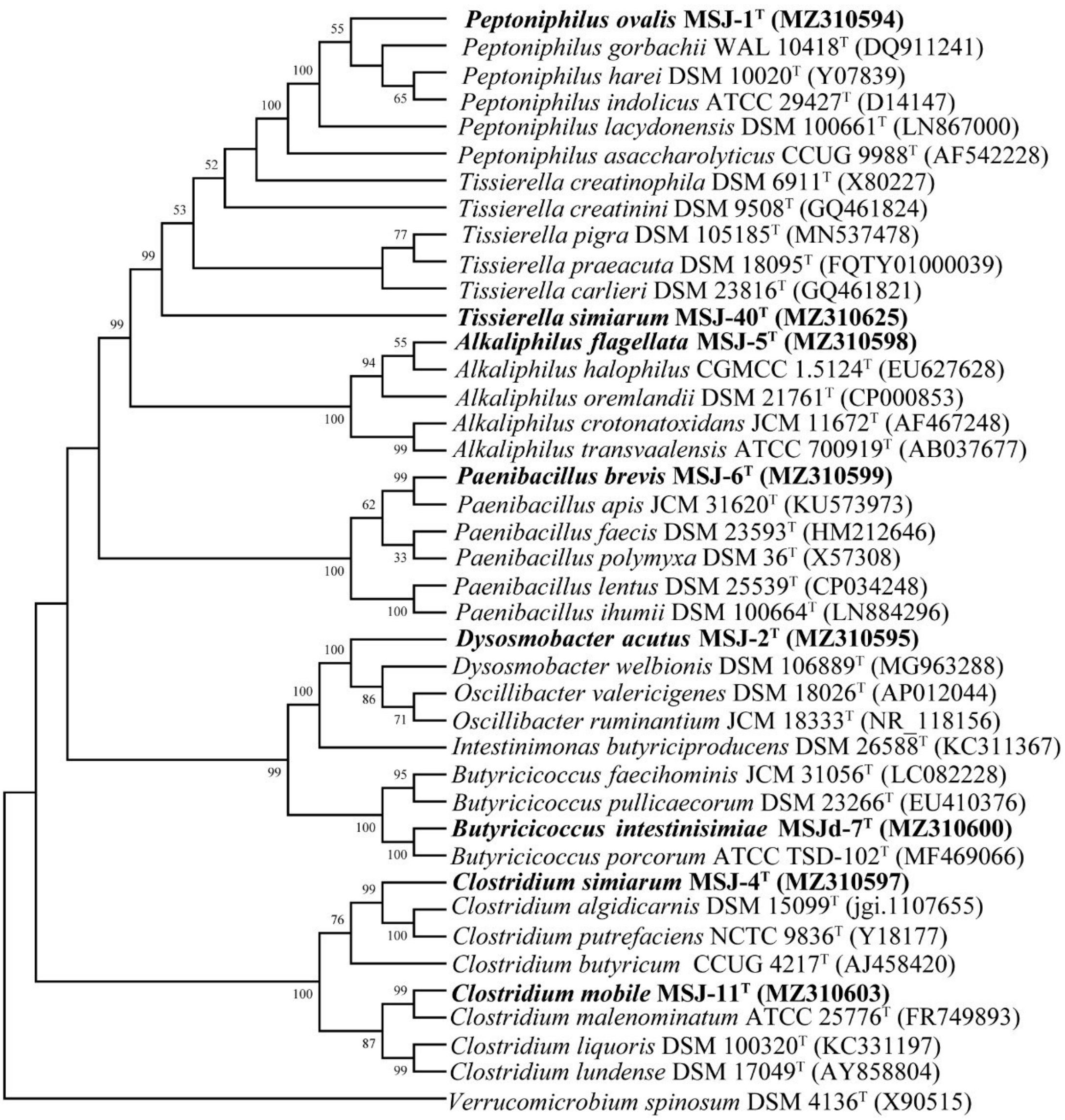


**Fig. S3 Maximum parsimony phylogenetic tree based on 16S rRNA gene sequences shows the relationship between nine strains and closely related microorganisms.**

Bootstrap percentages (>50%) based on 1, 000 replicates are shown at the nodes. GenBank accession numbers are given in parentheses. *Verrucomicrobium spinosum* DSM 4136$^T$ (X90515) was used as outgroup.
